# Causal Associations Between Body Mass Index and Mental Health: A Mendelian Randomization Study

**DOI:** 10.1101/168690

**Authors:** Nina van den Broek, Jorien L. Treur, Junilla K. Larsen, Maaike Verhagen, Karin J. H. Verweij, Jacqueline M. Vink

## Abstract

**Background:** Body Mass Index (BMI) is negatively correlated with subjective well-being and positively correlated with depressive symptoms. Whether these associations reflect causal effects or confounding is unclear.

**Methods:** We examined causal effects between BMI and subjective well-being/depressive symptoms with bi-directional, two-sample Mendelian randomization using summary-level data from large genome-wide association studies. Genetic variants robustly related to the exposure variable acted as instrumental variable (two thresholds were used; *p*<5e-08 and *p*<1e-05). These ‘instruments’ were then associated with the outcome variable. Pleiotropy was corrected for by two sensitivity analyses.

**Results:** Substantial evidence was found for a causal effect of BMI on mental health, such that a higher BMI decreased subjective well-being and increased depressive symptoms. No consistent evidence was found for causality in the other direction.

**Conclusions:** This study provides support for a higher BMI causing poorer mental health. Further research should corroborate these findings and consider non-linear effects and sex differences.

## Introduction

Obesity and poor mental health are two of the most pressing public health problems in modern-day society (1, 2). Previous studies have suggested that Body Mass Index (BMI) and mental health are correlated, such that increased adiposity is related to lower subjective wellbeing (3, 4) and more depressive symptoms (5). It is still unclear, however, whether these associations are (partly) due to causal effects and if so, in which direction. Longitudinal studies find support for associations in both directions (6–8). This may suggest that there are reciprocal causal effects, and/or overlapping third variables, including environmental factors (e.g., socioeconomic adversity) and/or shared genetic risk factors.

Mendelian randomization (9), an instrumental variable analysis using genetic variants as instruments, has been proposed as a tool to avoid biases from confounding (10–12). Analogous to randomized controlled trials, genetic risk factors related to the hypothesized cause (the exposure) are treated as natural experiments to estimate the consequences of the exposure on the outcome (the causal effect). Given the random nature of transmission of genes from parents to offspring, genetic risk factors are unlikely to be affected by confounders. Additionally, since genetic variation is fixed at conception, associations between genetic variants and outcomes cannot be attributed to reverse causality. A few studies have assessed associations between BMI and mental health using one-sample MR (e.g., 13, 14, 15). These yielded inconsistent results, potentially due to lack of power because of small sample sizes (N 1,731 to 4,145). Very recently, Wray et al. (16) performed the largest genome-wide association (GWA) meta-analysis of major depressive disorder (MDD). They also conducted MR and found strong evidence for a causal increasing effect of BMI on MDD, but no indication of a causal effect in the other direction.

Conventional, i.e. one-sample, MR requires genetic as well as phenotypic information on both the exposure and the outcome in a single sample, thereby limiting the available data and thus sample size. With two-sample MR, summary-level data from separate GWA studies of the exposure and the outcome variable are utilized, providing access to larger (existing) samples and thus increasing power. Using two-sample MR, Hartwig et al. showed weak, but consistent evidence of a causal effect of BMI on MDD, but not on bipolar disorder and schizophrenia (17). Whether causal effects are also relevant to less severe mental health outcomes, i.e., depressive symptoms or lower levels of subjective well-being, and whether such effects are bi-directional, is yet to be determined. We add to the current literature by performing bi-directional, two-sample MR with BMI on the one hand and subjective wellbeing and depressive symptoms on the other.

## Methods

MR is a type of instrumental variable analysis that utilizes genetic information as an instrument to test causal effects (10–12). This approach minimizes distorting effects of confounding and reverse causality, given that three assumptions are met; 1) the genetic instrument is predictive of the exposure variable, 2) the genetic instrument is independent of confounders, 3) there is no pleiotropy (the genetic instrument does not affect the outcome variable, other than through its possible causal effect on the exposure variable). Here, we apply *two*-*sample* MR, which takes the genetic instrument from a GWA study on the exposure variable (*gene*-*exposure association*) and then identifies the same genetic instrument in a GWA study on the outcome variable (*gene-outcome association).* Under a causal effect of exposure on outcome, genetic variants that predict the exposure variable should, through the causal chain, also predict the outcome variable (18).

All analyses were conducted with the two-sample MR package of *MR*-*Base* in R statistical software (19). Causal effects were tested *bi-directional:* from BMI (summary statistics from GIANT consortium; n = 339,224; 20) to depressive symptoms and subjective well-being (summary statistics from SSGAC consortium; n = 298,420, excluding 23andMe sample; 21) and vice-versa. Two sets of analyses were performed: one where only genetic variants that exceeded the threshold for genome-wide significance (*p*<5e-08) were included and one with genetic variants that exceeded a less stringent threshold (*p*<1e-05). Before analysis, genetic variants were pruned (*r*^2^<0.001) and where necessary proxies were identified (*r*^2^≥0.80). When a genetic instrument consisted of a single genetic variant, the causal effect was estimated using the Wald ratio method, which is computed as the gene-outcome association divided by the gene-exposure association (standard errors approximated by the delta method) (12). In case the genetic instrument consisted of multiple genetic variants, inverse-variance weighted (IVW) linear regression was applied, which sums the ratio estimates of all variants in a weighted average formula (22). Two-sample MR also allows correcting for pleiotropy when the instrument consists of multiple SNPs. We applied two commonly used sensitivity analyses. First, the *weighted median approach*, which can provide a consistent estimate of a causal effect even in a situation where up to 50% of the weight comes from invalid instruments (23). Second, *MR*-*Egger regression*, which adapts Egger’s test for small study bias in meta-analyses in such a way that it corrects for weak instrument bias in polygenic instruments (22).

Since we tested four different causal associations, we assumed a multiple testing alpha level of 0.0125 (0.05/4).

## Results

We found consistent evidence for a causal effect of BMI on subjective well-being, such that a higher BMI was associated with lowered well-being (**Table 1**). Only the MR-Egger results survived correction for multiple testing, but we found negative effects of similar magnitude across all methods and *p*-value thresholds (*p*<0.05). Evidence for a causal effect was also found from BMI to depressive symptoms. This positive effect, indicating that higher BMI increases depressive symptom, was seen for both *p*-value thresholds with the standard IVW method and the weighted median method, but it was not significant with the MR-Egger regression method.

**Table 1.** Causal Effect Estimates of Body Mass Index (BMI) on Subjective Well-Being (SWB) and Depressive Symptoms (DEP)

| Exposure | Outcome | *p*-value threshold<br>genetic instrument | # SNPs | IVW<sup>a</sup> |  |  | Weighted median |  |  | MR-Egger |  |  |
| --- | --- | --- | --- | --- | --- | --- | --- | --- | --- | --- | --- | --- |
|  |  |  |  | beta | *SE* | *p* | beta | *SE* | *p* | beta | *SE* | *p* |
| BMI | SWB | < 5e-08 | 78 | **-0.05** | **0.02** | **.023** | **-0.05** | **0.02** | **.021** | **-0.13** | **0.05** | **.011\*** |
| BMI | SWB | < 1e-05 | 179 | **-0.03** | **0.01** | **.046** | **-0.04** | **0.02** | **.024** | **-0.10** | **0.03** | **.004\*** |
| BMI | DEP | < 5e-08 | 78 | **0.05** | **0.02** | **.012\*** | **0.08** | **0.03** | **.002\*** | 0.07 | 0.05 | .148 |
| BMI | DEP | < 1e-05 | 180 | **0.04** | **0.02** | **.006\*** | **0.07** | **0.02** | **.003\*** | 0.07 | 0.04 | .061 |

Statistically significant associations are printed in bold and associations surviving correction for multiple testing (alpha level of 0.0125 (0.05/4)) are denoted with an asterisk. <sup>a</sup>IVW = Inverse-variance weighted; SE = standard error of the beta.

There was no evidence for a causal effect in the opposite direction, from subjective well-being and depressive symptoms to BMI (**Table 2**). There was only one suggestive, positive finding from subjective well-being to BMI, but it did not survive multiple testing correction and this test included only one genetic variant (i.e., rs2075677 in the CSE1L gene). Under the less stringent *p*-value threshold, where 27 genetic variants were included and we could correct for pleiotropy, this effect was not observed.

**Table 2.**
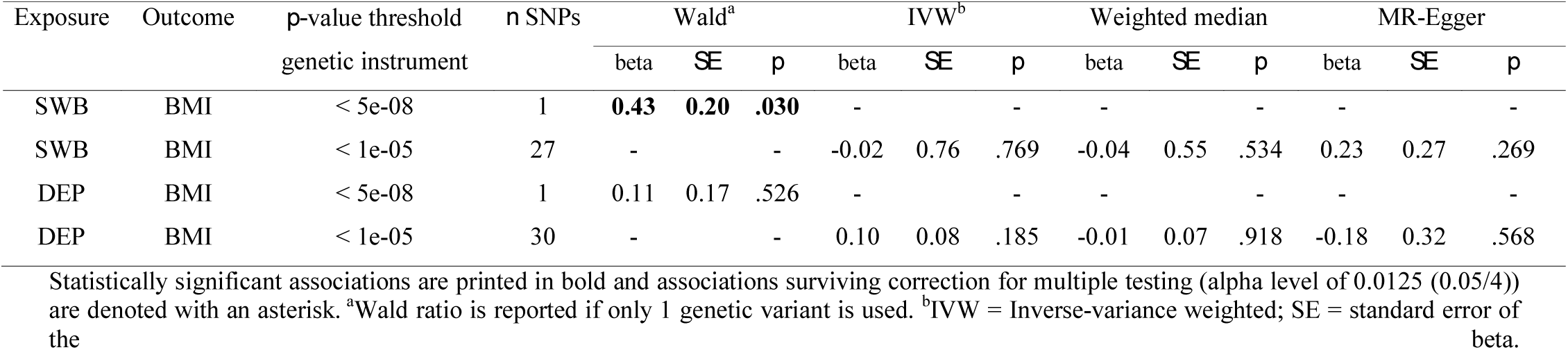
Causal Effect Estimates of Subjective Well-Being (SWB) and Depressive Symptoms (DEP) on Body Mass Index (BMI)

| Exposure | Outcome | $p$-value threshold | $n$ SNPs | Wald<sup>a</sup> | | | IVW<sup>b</sup> | | | Weighted median | | | MR-Egger | | |
| --- | --- | --- | --- | --- | --- | --- | --- | --- | --- | --- | --- | --- | --- | --- | --- |
| | | genetic instrument | | beta | SE | $p$ | beta | SE | $p$ | beta | SE | $p$ | beta | SE | $p$ |
| SWB | BMI | < 5e-08 | 1 | **0.43** | **0.20** | **.030** | - |  | - | - |  | - | - |  | - |
| SWB | BMI | < 1e-05 | 27 | - |  | - | -0.02 | 0.76 | .769 | -0.04 | 0.55 | .534 | 0.23 | 0.27 | .269 |
| DEP | BMI | < 5e-08 | 1 | 0.11 | 0.17 | .526 | - |  | - | - |  | - | - |  | - |
| DEP | BMI | < 1e-05 | 30 | - |  | - | 0.10 | 0.08 | .185 | -0.01 | 0.07 | .918 | -0.18 | 0.32 | .568 |

Statistically significant associations are printed in bold and associations surviving correction for multiple testing (alpha level of 0.0125 (0.05/4)) are denoted with an asterisk. <sup>a</sup>Wald ratio is reported if only 1 genetic variant is used. <sup>b</sup>IVW = Inverse-variance weighted; SE = standard error of the beta.

Forest plots depicting the effects of all individual genetic variants included in the genetic instruments are provided in the **Supplementary material**. We also performed ‘leave-on-out’ sensitivity analyses which entailed repeating the analyses of multi-SNP instruments after excluding each of the SNPs, one at a time. Results indicated that the causal effects were consistent and not driven by one or a few specific SNPs.

## Discussion

With bi-directional, two-sample Mendelian randomization, we found substantial evidence for a causal effect of BMI on mental health such that it decreased subjective well-being and increased depressive symptoms, but not the other way around. This is in line with a recent, two-sample MR study which reported weak evidence for a higher BMI being causally related to a higher risk of MDD (15). It also concurs with a large bi-directional, one-sample MR study that reported evidence for a causal effect of BMI on MDD, but not in the other direction (16), and with two smaller one-sample MR studies that reported causal effects of BMI on common mental disorders in men (but not in women) (13) and on depressive symptoms (15). In contrast, two other one-sample MR studies did not find evidence for a causal effect of BMI on MDD or on a depression score based on a variety of assessments of depression diagnosis or symptoms (14, 24). These contradictory findings may be explained by heterogeneity between the outcome variables, a lack of power in the smaller, one-sample MR studies, or it may be the result of violations of the assumptions of the MR design (25). The findings in the present study are strengthened by the two-sample MR approach, which has considerably higher power to pick up on causal effects compared to the one-sample MR approach, and allows sensitivity analyses to correct for pleiotropy.

As for the mechanisms underlying causal effects, it has been reported that weight-related problems such as physical illness and weight stigmatization negatively affect mental health (26). Also, a higher BMI could lead to inflammation and a dysregulated hypothalamic-pituitary-adrenal (HPA)-axis, which in turn could induce the development of depressive symptoms (27). The exact causal mechanism between BMI and mental health is a topic that needs further investigation (26, 28).

While two-sample MR greatly improves power compared to one-sample MR (29), our genetic instruments still only account for a very small part of the variance. Replicating this study when even bigger GWA studies become available is warranted. Another limitation is that we were unable to address any non-linear (causal) effects. It seems plausible that the association between BMI and mental health is inverse u-shaped, such that both very low and very high levels of BMI lead to impaired mental health (3, 30). In addition, it would be interesting to look at sex differences given that there is evidence that weight stigma has worse consequences for obese women than for obese men, and that the association between mental health and obesity can also differ between the sexes (31).

To our knowledge, this is the first study to bi-directionally assess causal associations between BMI and mental health. Our findings support the idea that a higher BMI causally leads to a decrease in mental health. Future research needs to corroborate our findings, ideally with stronger genetic instruments and whilst also considering non-linear effects, and possible sex differences.

## Acknowledgments

We are thankful to investigators of GIANT and SSGAC for making their summary statistics publicly available.

